# Modelling forest dynamics using integral projection models (IPMs) and repeat LiDAR

**DOI:** 10.1101/2025.01.06.631514

**Authors:** Alice Rosen, Robin Battison, Christina M. Hernández, Oliver Spacey, Jessie McLean, Suzanne Prober, Samuel Gascoigne, Sean McMahon, Tommaso Jucker, Roberto Salguero-Gómez

**Affiliations:** Department of Biology, University of Oxford, South Parks Road, Oxford, OX1 3RB, UK; School of Biological Sciences, University of Bristol, 24 Tyndall Avenue, Bristol, BS8 1TQ, UK; CSIRO Centre for Environment and Life Sciences, Underwood Avenue, Floreat, Western Australia; Smithsonian Environmental Research Center, 647 Contees Wharf Road, Edgewater, MD 21037-0028, USA

**Keywords:** Airborne Laser Scanning (ALS, or LiDAR), demography, forests, Integral Projection Model (IPM), remote sensing, structured population model

## Abstract

1. Estimating the life histories and population dynamics of trees is important for predicting how forests respond to climate change and disturbances. To do so, ecologist often use stage-structured population models, which explicitly account for individual heterogeneity. However, the data needed to parameterise these models is typically constrained by time- and labour-intensive field methods.
2. Despite growing access to remote sensing imagery that could potentially ‘satiate’ these data-hungry population models, these data remain underutilised in population ecology. Here, we demonstrate a pipeline that integrates repeat-LiDAR data with a type of structured demographic model (Integral Projection Model, IPM) to examine forest-wide dynamics in response to environmental drivers. Using Australia’s Great Western Woodlands as a case study, we model the survival and growth of ∼40,000 eucalypt trees over a decade to estimate their life expectancy and characterise multiple stages of growth (height growth and crown expansion).
3. Our results indicate distinct responses of small trees (where vital rates are predicted by height) and large trees (which invested predominantly in expanding crown area) to proxies for competition and soil moisture (local canopy density and topographic wetness index, respectively). Parameterising structured population models using LiDAR data therefore offers a step-change in perspective towards more ecologically meaningful forest dynamics.
4. To broaden the application of this pipeline, we highlight three priorities: (1) Application to more complex systems such as mixed species stands and dense, multilayered canopies); (2) Incorporating complete life histories, including reproduction and early life stages; and (3) Using in long-term or comparative studies through improved availability of high-quality, comparable, repeat-LiDAR surveys. By combining remote sensing data with detailed insights from field-based studies, this pipeline represents a scalable framework for guiding forest management and conservation decisions.

## 1 | INTRODUCTION

The world’s forests are facing growing pressure from climate change and land-use intensification, with major implications for carbon and water cycling, biodiversity, and people (McDowell et al., 2020; Seidl & Turner, 2022). However, projecting how forests will respond to these changes is inherently challenging due to the vast spatial and temporal scales at which forests operate. In response, a wide range of modelling approaches have been developed to understand and predict the impact of these threats (Decocq et al., 2023).

Demographic models allow us to take into account the varied life history strategies of trees – the schedules of the vital rates of survival, growth and reproduction (Clark & Clark, 1992), and estimate key life history traits (*e.g.,* generation time, mean life expectancy; Bienvenu et al., 2015; Salguero-Gómez et al., 2016) and their sensitivity to changing conditions (Franco & Silvertown, 2004; Werner & Peacock, 2019). These models therefore allow us to examine how specific threats, such as drought, may disproportionately impact the growth of, for example, the largest trees (Bennett et al., 2015; Stovall et al., 2019; Trouillier et al., 2019), while threats such as seed harvesting may instead limit juvenile recruitment (Peres et al., 2003).

Demographic approaches are, however, data hungry (Doak et al., 2005; Ramula et al., 2009), requiring data on the vital rates for hundreds – if not thousands – of individuals in the population of interest. For long-lived trees, it is rarely possible to observe an individual from the beginning to the end of its life (Munné-Bosch, 2020). Instead, vital rates can be estimated by monitoring individuals in a population at two or more time points (Rees et al., 2014). However, producing robust estimates of vital rates is still easier said than done. In particular, obtaining data on tree vital rates at the beginning (*i.e.*, reproduction/recruitment; Bogdziewicz et al., 2024) and end of their life cycle (*i.e.*, adult mortality; McMahon et al., 2019) can be notoriously challenging (Ruiz-Benito et al., 2020).

Although tree growth data are more abundant than recruitment or mortality data (Ohse et al., 2023), they primarily focus on changes in diameter at breast height (DBH) of the main trunk (Lines et al., 2022) – a simple and widely used metric linked to aboveground biomass (Chave et al., 2014). While valuable, DBH alone does not capture how trees adapt and shift their above-ground growth allocation strategies over their life cycle, including their investment in vertical and lateral crown expansion (King, 2011; Pretzsch et al., 2013). Moreover, most demographic data for trees comes from well-studied permanent plots (*e.g.,* Yang et al., 2018), forest inventory data (*e.g.,* Kunstler et al., 2021), or tree-ring studies (*e.g.,* Brienen & Zuidema, 2006). As well as being time, cost, and labour intensive, these methods necessarily compromise on the spatial scale and/or the number of individuals used to represent entire populations (Ohse et al., 2023; Ruiz-Benito et al., 2020).

Remote sensing, particularly LiDAR (Light Detection and Ranging), has unlocked new ways of studying forests at scales relevant for informing policy and management decisions (Camarretta et al., 2020; Masek et al., 2015). LiDAR sensors emit high frequency laser pulses and measure the time it takes for those pulses to return to the sensor. Mounted on an airplane or unoccupied aerial vehicle (UAV), LiDAR therefore allows us to capture the 3D structure of forest canopies at scale, in the form of a detailed point cloud model (Camarretta et al., 2020). As a result, this technology also provides a change in perspective – shifting our focus away from traditional trunk-based measures like DBH (where the majority of the tissue is dead; Magel et al., 1994) and towards the canopy, where biologically relevant processes and dynamics take place (Lines et al., 2022).

LiDAR data are becoming more widely and freely available (Beland et al., 2019), offering the opportunity to extend the application of demographic modelling approaches across broader scales and forest types. Where options for accessing these data were previously cost-prohibitive, more affordable platforms for collecting LiDAR data (*e.g.,* UAVs), combined with initiatives for open data sharing (Kampe et al., 2010), have improved access to large-scale, high-resolution datasets. In addition, dedicated software (*e.g.,* CloudCompare; https://danielgm.net/cc), R packages (Atkins et al., 2022; Roussel et al., 2020) and robust methodologies for LiDAR processing and analysis (Fischer et al., 2024) have facilitated the use of LiDAR data in ecological studies, even by non-experts. Despite the quantity of remote sensing data now being generated, and an uptake in the use of LiDAR data by ecologists in recent years, these datasets remain under-utilised in demographic modelling.

Here, we introduce a pipeline for combining remote sensing data with a structured demographic model to demonstrate their combined potential for understanding forest dynamics at scale. As a case study, we model the demographic changes of ∼40,000 Eucalypt trees in a fast-drying region of southwestern Australia (Brouwers & Coops, 2016; CSIRO, 2022) and assess how the survival, growth, and remaining life expectancy of individuals in this population are affected by different biotic (canopy density) and abiotic conditions (soil moisture). Then, we discuss the advantages and limitations of a remote-sensing approach to forest demographic modelling. Finally, we highlight three areas of future research that are key to broadening the application of this framework: to more complex systems (*e.g.,* mixed species stands and dense-multilayered canopies); across complete life histories (*e.g.,* to include reproduction and early life stages); and for long-term or comparative studies (*e.g.,* through improved availability of high-quality, repeat-LiDAR surveys and integration with field data).

## 2 | METHODS

### 2.1 | Integral Projection Models (IPMs)

The heterogeneity of individuals in most populations is key for understanding the ecological processes that shape their viability (Ohse et al., 2023). These processes include how different individuals in the population contribute to successional dynamics (Jansen et al., 2012), determine the overall resilience of the ecosystem (Guyennon et al., 2023), and differentially respond to disturbances (Zambrano & Salguero-Gómez, 2014). Indeed, a relatively small number of ‘super-performing’ individuals with persistently fast growth rates (perhaps due to an environmental and/or genetic advantage) may ultimately shape the population dynamics of long-lived species (Jansen et al., 2012). Stage-structured models are ideally placed to incorporate this individual heterogeneity to understand and predict the performance of natural populations. Within stage-structured models, integral projection models (IPMs) (Coulson, 2012; Easterling et al., 2000; Merow et al., 2014) explicitly examine how continuous traits (*e.g.*, size) that characterise individuals in the population relate to their vital rates of survival, growth, and reproduction. By using one or more continuous traits (Zambrano & Salguero-Gómez, 2014) an IPM allows us to create population projections that identify the individuals most at risk from threats such as climate change and the impact on the population as a whole (Laurans et al., 2024; Ohse et al., 2023).

Here, we use a case study to demonstrate the pipeline for parameterising an IPM using remotely sensed data (Fig. 1). The pipeline broadly involves five key steps: 1) Using repeat LiDAR data to track individuals through time; 2) Selecting relevant traits (*e.g.*, tree height and crown area) for predicting vital rates; 3) Modelling vital rates from the selected traits, as well as relevant LiDAR-derived environmental drivers; 4) Building a multi-stage IPM that captures the primary (height) and secondary (crown expansion) growth of these trees; and 5) Analysing the IPM to understand the relative influence of biotic (*e.g.,* density dependence) and abiotic drivers (*e.g.*, soil moisture) on key emerging properties of the population model, such as life expectancy.

**Figure 1.**
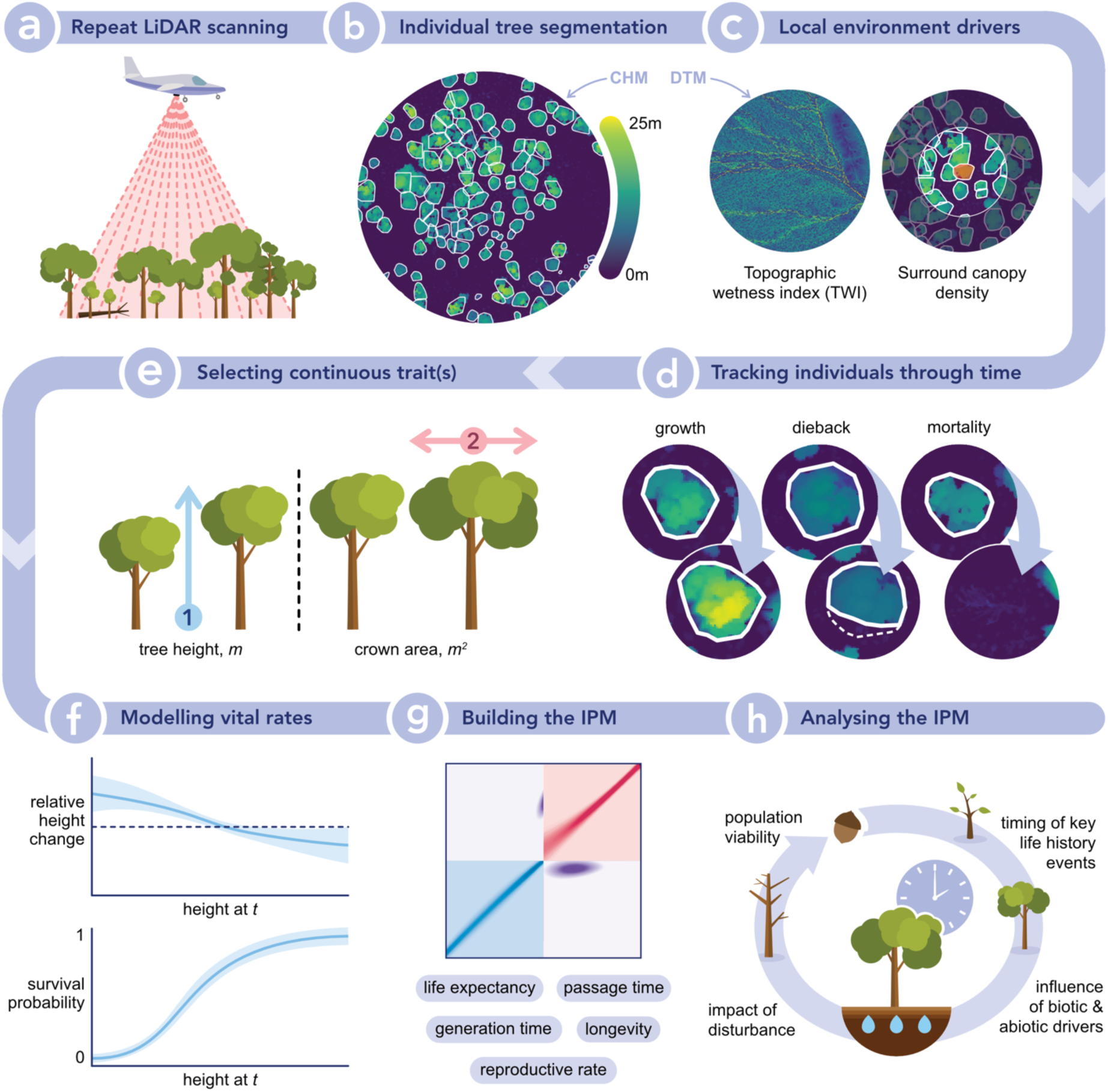
Pipeline for combining LiDAR data with structured demographic models for ecological inference. **a**) The population is surveyed at appropriate time intervals with a LiDAR scanner to produce high-resolution 3D point clouds. The point clouds are processed to produce canopy height models (CHMs) and digital terrain models (DTMs). **b**) Individual tree segmentation is performed using a suitable algorithm designed to work directly on the CHM (*e.g.,* Dalponte & Coomes, 2016). **c**) Relevant environmental variables can be derived. *E.g.,* topographic wetness index (TWI) is an indicator of soil moisture accumulation (Kopecký et al., 2021) and is calculated from the DTM. Local competition metrics can also be inferred from the canopy density surrounding a focal tree, calculated using the CHM. **d**) It is then necessary to match individuals across time points using an appropriate crown-matching algorithm (Battison et al., 2024; Olsoy et al., 2024), before measuring changes in size and classifying mortalities. **e**) Relevant continuous traits (*e.g.,* height and crown area) are explored and selected based on their ability to predict survival and growth across the life cycle. **f**) The selected traits and environmental parameters are used to model the vital rates using regressions. **g**) The integral projection model (IPM) is constructed based on the selected vital rate models, and key life history traits (*e.g.,* mean life expectancy, longevity, generation time) are calculated. **h**) Finally, analysis of the IPM and its predicted life history traits are used to test important ecological questions.

### 2.2 | Case study: Eucalypt woodlands of southwestern Australia

#### Study site and species

The Great Western Woodlands Terrestrial Ecosystem Research Network (TERN) SuperSite (30°11’ S, 120°39’ E, 430-475 m a.s.l) is located in the Great Western Woodlands of south-western Australia, with key goals including detecting temporal ecological change and informing ecological management in response to fire and climate change (Prober et al., 2023). The Great Western Woodlands is a semi-arid region, comprising one of the largest intact temperate woodlands in the world (Watson et al., 2008). The region experiences a seasonal climate with a prolonged dry season, mean annual rainfall of 265 mm, and a mean annual temperature of 19°C. Importantly, due to climate change, increases in temperature and higher drought frequency are projected in the region over the next two decades (CSIRO, 2022; Peters et al., 2021; Prober et al., 2012).

Increasingly warm, dry conditions in the region are contributing to more frequent wildfires (Prober et al., 2012). Indeed, the primary driver of vegetation dynamics in the region is stand-replacing fires. An estimated 38.8% of woodland area in the Great Western Woodlands has burned at least once in the past 50 years (Jucker et al., 2023). However, the study site, a 25 km^2^ area at the core of the SuperSite, is predominantly old-growth woodland that has not been disturbed by wildfires for hundreds of years. The site is largely dominated by four eucalyptus species: *Eucalyptus salmonophloia* (the most abundant), as well as *E. salubris*, *E. transcontinentalis*, and *E. clelandiorum*. These four species share a common ecological strategy: they are obligate-seeders that depend on their canopy-held seed bank to regenerate after fire (Gosper et al., 2018). The obligate seeders lack thick bark or the ability to resprout, so even low severity fires result in tree mortality followed by dense seedling stands. In the absence of fire, seedling recruitment rates are substantially lower, but the ecosystem can persist through gap phase recruitment (Gosper et al., 2018). In this old-growth phase, the woodland is characterised by a relatively sparse, open canopy with large, single-stemmed trees. The four dominant species can grow to be 10-25 m in height, and live to at least 400 years (Jucker et al., 2023). Mulga scrub (*Acacia* sp.) and heathland are also present as mosaics with eucalypt woodlands in the Great Western Woodlands landsacpes, but these account for <0.1% of the study site, and so they were excluded from our analyses.

#### LiDAR data

Airborne LiDAR data were acquired in May 2012 and May 2021 across our 5 × 5 km^2^ study site, producing a 3D “point cloud” of the woodland. The 2012 LiDAR survey was conducted by Airborne Research Australia (https://www.airborneresearch.org.au) using a motorised glider (Diamond Aircraft, HK36 TTC-ECO) flown at 300 m altitude and mounted with a RIEGL LMS-Q560 scanner. The 2021 data were collected by Aerometrex (https://aerometrex.com.au) using a fixed wing aircraft (Cessna 404 Titan), flown at 1,100 m altitude and mounted with a REIGL VQ-780ii scanner. Both surveys achieved similar, high pulse densities, despite differences in the acquisition parameters and instrumentation (21.4 m^-2^ and 23.6 m^-2^ for the 2012 and 2021 surveys, respectively).

From these high-quality 3D point clouds, we generated 2D maps of canopy height, known as canopy height models (CHMs; Fischer et al., 2024). The georeferenced point clouds for both LiDAR surveys were processed using a combination of CloudCompare (https://danielgm.net/cc), QGIS (https://qgis.org) and R version 4.1.0 (R Core Team, 2021). See supplementary information for details. We used the pit-free algorithm in the *lidR* R package (Khosravipour et al., 2014) to produce a 0.5 m resolution normalised canopy height model and a 5 m resolution digital terrain model (DTM). We selected this algorithm to optimise comparability between the two surveys, as it is robust when applied to high-quality point clouds (Fischer et al., 2024; Zhang et al., 2024). All subsequent data processing and analyses were conducted in R version 4.1.0.

### 2.3 | Using LiDAR to track individuals through time

#### Individual tree segmentation

To estimate individual tree survival and growth, the first step is to identify and outline their position within the LiDAR dataset. Many tree detection algorithms are available for this task. Some of these algorithms work directly on the canopy height model (CHM; *e.g.,* Dalponte & Coomes, 2016; Cao et al., 2023), while others work on the underlying point cloud (*e.g.,* Yang, Kang, et al., 2019). As well as being computationally cheaper, CHM-based algorithms are generally more robust to differences in how the data are acquired (Fischer et al., 2024; Mielcarek et al., 2018). To segment individual trees, we applied the widely-used *dalponte2016* crown delineation algorithm (Dalponte & Coomes 2016), following the process outlined in Battison et al. (2024). We selected this algorithm as it outperformed alternative algorithms in terms of its ability to accurately detect trees and to estimate their crown sizes when compared to a subset of manually delineated tree crowns (Battison et al., 2024). Briefly, this algorithm works directly on the canopy height model, applying a local maximum filter to locate the tallest points of individual trees before finding the borders of their crowns. A local maximum filter is applied within a moving window that is allowed to vary in size with the local height of the canopy. Window size is based on the allometric constraints between tree height (m) and crown area (m^2^), which we determined from the subset (n = 797) of manually delineated trees. Alternatively, tree allometry databases such as Tallo (Jucker et al., 2022) can be used to model the appropriate window size.

#### Local environment drivers

With the tree crowns delineated, we used the same LiDAR datasets to gain additional context about the local environment of individual trees. In this case study, we explored an example of a biotic and abiotic driver (proxies for density dependence and drought) to investigate how these factors may drive variation in vital rates and key emerging properties of the population. First, as an indicator of above- and belowground competition, we estimated the neighbourhood canopy density for each delineated tree (Vanderwel et al., 2020). We calculated canopy density as the mean height of the 2012 canopy height model within a 25 m radius around the perimeter of each tree crown (Battison et al., 2024; Zhao et al., 2006). Then, as a proxy for local soil moisture and nutrient availability, we calculated the topographic wetness index (TWI). TWI is a widely used metric that describes how water is likely to accumulate across the landscape based on topographic factors such as the underlying slope and terrain (Kopecký et al., 2021). We measured TWI from the 5 m resolution digital terrain model using the *dynatopmodel* package in R. We assigned each tree a value of TWI based on its coordinates. Later, we investigate the effect of these parameters within our vital rate models to understand how they may impact survival and growth.

#### Crown matching across time

To track changes in tree size between the two LiDAR surveys, we matched segmented tree crowns between 2012 and 2021 using the approach developed by Battison et al. (2024). First, we removed individuals whose centroid in 2021 fell outside of the crown boundary in 2012. Then, we filtered out instances where more than one centroid was found within the boundary of a single crown at the other time point. This step aimed to minimise over-segmentation, whereby individual trees are mistakenly delineated into separate, smaller individuals. Retaining these individuals would introduce extreme height and crown area changes between the two time points. Finally, although the four eucalypt species make up most of the community in the study site, there are also some smaller shrubs present. We therefore removed crowns smaller than 3 m in height or with a crown area of less than 9 m^2^ in 2012 to ensure these shrubs were filtered out. The outcome of this filtering process was a subset of time-matched eucalypt tree crowns (n = 39,286) that we used to model tree growth in the construction of the IPM.

### 2.4 | Estimating vital rates from LiDAR

#### Selecting relevant traits

We next considered which continuous traits might be suitable for modelling vital rates such as survival and growth. To do so, we explored size-related traits that can be easily derived from LiDAR data: tree height, crown area, and the interaction between height and crown area (which we will refer to as ‘tree size’) to capture allometric scaling relationships (Lines et al., 2022). To guide our decision, we investigated the direction of tree growth in terms of height, crown area and overall tree size change across the full distribution of tree sizes (see supplementary information). In doing so, we observed that smaller trees prioritise height growth, whereas larger trees switch instead to prioritising crown expansion, a common strategy observed among trees (Franceschini et al., 2016; King, 2011). Although initial tree size (in 2012) was a strong predictor of tree size in 2021 (*R^2^* = 0.96; *p* < 0.001), this trait hides the changing trajectory of how trees allocate their resources to growth when they are small or large – towards height growth and crown expansion, respectively. We therefore decided to use two continuous traits: height to model the vital rates of small trees, and crown area to model the vital rates of large trees (details below).

#### Modelling vital rates

To capture how vital rates may differ between these two stages of development (small and large trees) we modelled vital rates separately for each size class. We chose an appropriate cut-off point to differentiate between small and large individuals (Fig S2), and then fit separate vital rate functions to each discrete category. See supplementary information for details on how the size cut-off was selected. Briefly, we chose a cut-off value based on the point at which relative height change and relative crown area change were predicted to be equal (tree height = 7.27 m). This value was close to the median value of tree height in 2012 (7.06 m). We note that there are many ways to determine a suitable cut-off between multiple possible discrete categories in an IPM, and this choice should be based on knowledge of the system. Individuals smaller than the cut-off in time *t* (2012) and *t* + 1 (2021) were assigned to the ‘minor class’, whilst trees larger than that cut-off in *t* and *t* + 1 were assigned to the ‘major class’ (Fig. 2a). Additionally, we modelled the growth of trees that transitioned over the cut-off from *t* to *t* + 1 (‘promotion’), and those that fell below it (‘demotion’). Trees in the minor class were modelled by a continuous measure of (log-transformed) height, whereas those in the major class were modelled by a continuous measure of (log-transformed) crown area.

**Figure 2.**
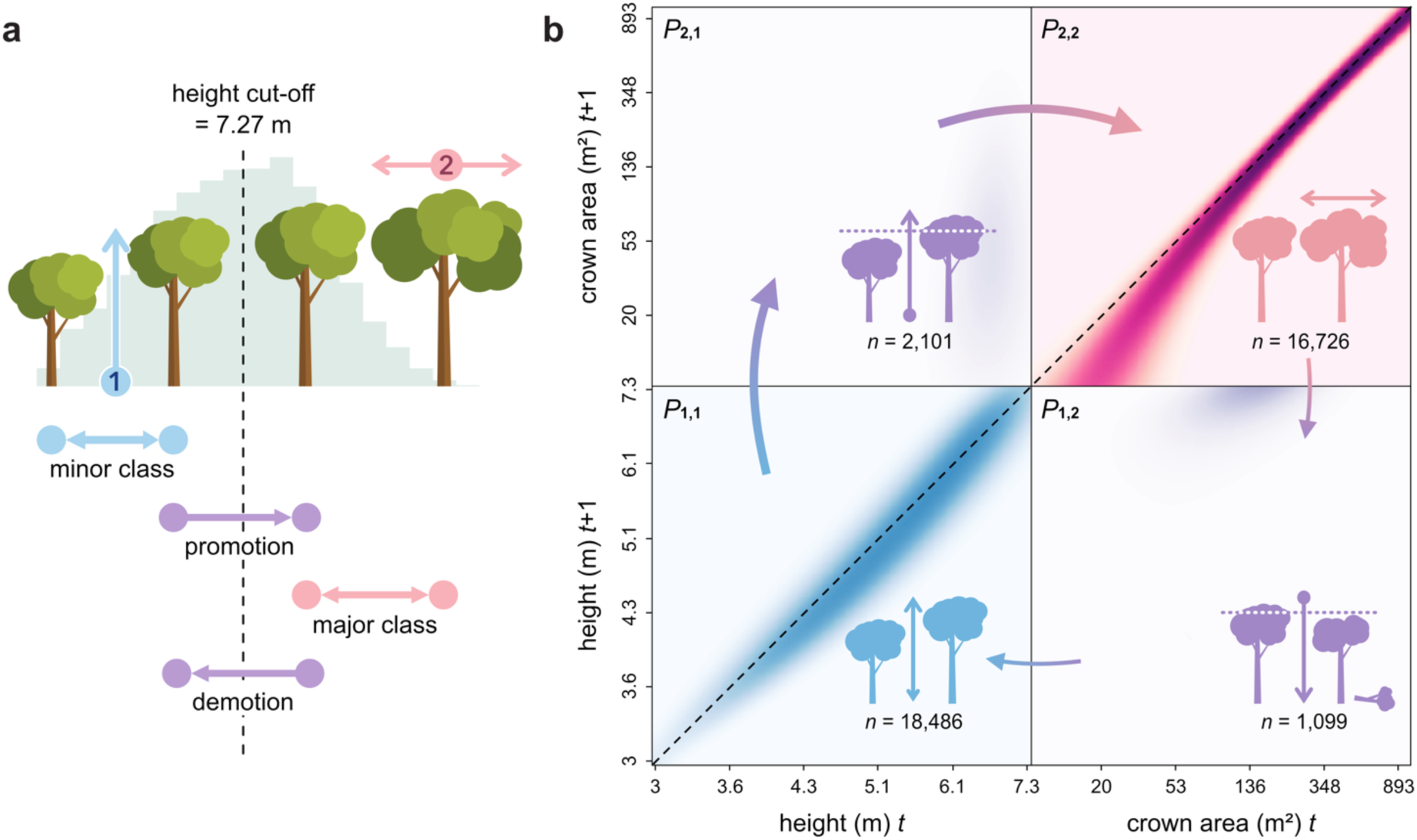
A two-stage integral projection model (IPM) used to explore population dynamics of a cohort of eucalypts in southwestern Australia. **a**) Trees were assigned to a class based on their size at time *t* and *t* + 1, relative to a height threshold of 7.27 m. **b**) The *P*-kernel (survival [Eq. 1] and growth [Eq. 2] component of an IPM) shows the probability of a tree of size *x* at time *t* surviving and growing to be size *y* at time *t* + 1, where colour intensity indicates probability. Cell *P*_1,1_ (blue) represents trees in the minor class (based on height); *P*_2,2_ (pink) represents those in the major class (crown area); and *P*_2,1_ and *P*_1,2_ (purple) represent transitions between the two main classes – promotion and demotion, respectively. The clockwise arrows represent the theoretical transition of a tree through its lifetime as it grows, surpasses the height cut-off, and then shrinks again – perhaps due to dieback or because it has lost some of its branches after a storm.

To predict tree survival and growth in each of the discrete size classes, we compared a set of ecologically meaningful models. In each instance, we explored the contribution of soil moisture (topographic wetness index, TWI) and local competition (canopy density) to vital rates. Selection of the best supported model was based on a combination of the Akaike information criterion (AIC) combined with our biological knowledge of the system.

#### Survival

To model the survival of individual trees, we first classified mortality events via a two-step process. First, in the matched crowns subset (n = 39,286; described in section 2.3), trees that experienced >30% reduction in height and/or crown area between the two surveys were classified as dead, following Duncanson & Dubayah (2018) and Ma et al. (2023). Second, to capture mortality events where crowns could no longer be detected in the second survey, we overlayed the polygons of 2012 delineated trees onto the 2021 CHM, which allowed us to measure the height and crown area change of unmatched trees. Then, to classify tree death, we applied the same >30% reduction threshold as above. We performed a sensitivity analysis to examine how the choice of percentage threshold shapes mortality rates (Table SX). This approach is recommended to assess the robustness of tree mortality estimates, given the variety of ways that dead trees may be detected in the data.

For each of the discrete classes (minor and major), we modelled the probability that an individual tree survives from time *t* to *t* + 1, given its size at time *t*, using logistic regressions. First, we compared a linear model of size against a quadratic model, which can accommodate a decline in vitality at large sizes (Coomes & Allen, 2007). Then, these models were compared against more complex models that additionally accounted for TWI alone, canopy density alone, or the additive effects of TWI and canopy density together to test their effect on a population predicted to experience rapid warming and drying (CSIRO, 2022). The selected model for both the minor and major classes was a generalised linear model with a logit link function, with the form:

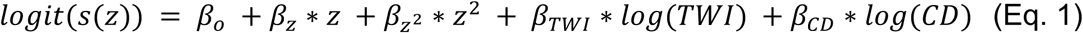

where z is the log-transformed size of an individual at time *t*, TWI is topographic wetness index, CD is canopy density, and the *β*’s are the fitted coefficients.

It is worth noting that, in this model, the probability of survival (*s(z)*) only depends on size at time *t*, regardless of whether individuals remain in the same class, or promote/demote to another (Fig. 2). More complex models that evaluate vital rate trade-offs are described elsewhere (Miller et al., 2012).

#### Growth

To model the growth of individual trees, we quantified changes in tree height and crown area between 2012 and 2021. To do this, we again used the 39,286 matched tree crowns described above, as well as the CHMs from 2012 and 2021 (Fig. 1). We defined tree height as the mean value of the canopy height model within the delineated tree crown. The mean height of tree crowns is typically less affected than the maximum value to outliers resulting from artefacts such as clouds. In this case, however, the mean value was highly correlated with the maximum value (*R*^2^ = 0.87; *p* < 0.001), indicating that maximum height was mostly unaffected by artefacts. Crown area was defined as the area of the polygon that represents the delineated crown. Both size measures were calculated using the *terra* package in R (Hijmans, 2024).

Next, we modelled the expected distribution of tree sizes at time *t* + 1, for an individual of size *x* at time *t*, conditional on having survived from time *t* to *t* + 1. The overall distribution of tree sizes at time *t* + 1 is then obtained by summing the resulting distributions over all observed tree sizes. We performed ordinary least-squares regressions for the response variables (height or crown area in time *t* + 1) against their size in time *t*. We selected a growth function that models expected size at time *t* + 1 using a quadratic regression of size at time *t* (except for the demotion class, where a linear growth function was selected instead; Table SX), and additionally accounts for log-transformed TWI and canopy density:

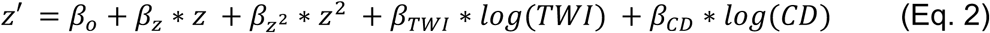

where z’ is the log-transformed size of an individual at time *t* + 1 and all other variables are as previously defined. Tree size at time *t* may also influence the variance in size at time *t* + 1. Therefore, to account for the variation in individual growth trajectories, we estimated the size-dependence of this variance by regressing the residuals in the growth model against size at time *t*. The selected model for the residuals was quadratic for all classes except the demotion class, where the most parsimonious model was a simple linear model (Table S1).

### 2.5 | Building the IPM

To model the demographic changes of the study eucalypt woodland in the context of a changing climate, we parameterised an integral projection model (IPM) using LiDAR data described above. IPMs are demographic models that describe population dynamics across discrete time, where individual heterogeneity is accommodated by assigning individuals in the population to values along one (or multiple; Fig. 2) continuous trait(s) (*e.g.*, height, crown area; Zambrano & Salguero-Gómez, 2014). In this case study, we focus on the survival and growth of the existing population under different conditions of canopy density and soil moisture. These vital rates allow us to investigate multiple life history traits, such as remaining life expectancy, that have implications for short-term carbon sequestration (Needham et al., 2022). The component of an IPM that captures the survival and growth transitions of an existing cohort (without reproduction) is known as the *P*-kernel:

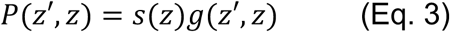

where *s*(*z*) is the probability that an individual of trait value *z* (here height or canopy area) at time *t* (2012) will survive to *t* + 1 (2021) (Eq. 1), and *g*(*z*′, *z*) is the growth function that describes the probability density of size *z*′ at time *t* + 1, given an individual’s size *z* at time *t*, conditional on having survived that time period (Eq. 2). We note that to model the reproductive component (typically referred to as the *F*(*y*, *x*) kernel), one would also need data on fecundity rates and establishment probabilities (*e.g.,* Easterling et al., 2000). See applications of our framework to systems with reproductive data in section 4.2 ‘Completing the life cycle’.

#### Constructing a two-stage IPM

To account for the changing growth trajectory of the eucalypt individuals (from height growth to crown expansion), we constructed a two-stage *P*-kernel. Our approach is similar to that introduced by Zambrano & Salguero-Gómez (2014) used to examine the population dynamics of a tropical tree in response to forest fragmentation based on the changes in height of saplings and changes in crown area of adults. To accommodate the two discrete classes in our population model (minor and major, Fig. 2), we built a 2 × 2 mega-matrix ***M*** (Goodman, 1968). Each of the four emerging cells in the matrix then accommodates vital rates as a function of continuous traits (height and/or crown area). The four cells, as introduced in Figure 2, correspond to *P*_1,1_: survival and growth of minor class individuals (based on height) remaining as such; *P*_2,1_: promotion of individuals in the minor class (measured as a function of height) to the major class (crown area); *P*_2,2_: survival and growth of major class individuals (crown area) remaining as such; and *P*_1,2_: dieback from major (crown area) to minor (height) class individuals. While in this case study we explore a life history trait based on growth and survival, data on reproduction (the *F-*kernel described above) can also be accommodated in the cell ***M***_1,2_, together with *P*_1,2_. We modified the form of the above *P*-kernel to include the probability that an individual of size *z* at time *t* will remain within the same size class at *t* + 1 (as opposed to being promoted or demoted from the current size class):

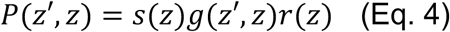

We used a generalised linear model with logit link function to model the probability of a tree remaining in its size class from time *t* to *t* + 1 (*i.e.,* the tree is measured by the same continuous state variable – height or crown area – from one time to the next). The inverse probability therefore describes the probability of a tree in the minor class being promoted, or a tree in the major class being demoted. The selected model for the probability of trees remaining in their current class is:

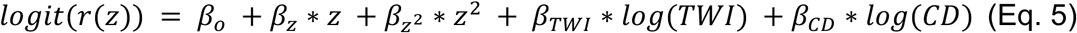

Sometimes, the predicted values for size-specific survival may fall outside of the size range of the kernel. This feature of IPMs causes individuals to be ‘evicted’ from the model, leading to unrealistic population projections (Ellner et al., 2016; Merow et al., 2014). Eviction is a common problem that can result from several factors, including the choice of growth model, or the selected size limits of the kernel. In this case, we corrected for eviction by forcing the column sums of each *P*-kernel to equal the predicted values of size-specific survival that emerge from the survival and remain functions described above. For the construction of the IPM, we used eviction-corrected kernels for all but the major class kernel. Instead, we used the uncorrected major class kernel to retain the natural reduction in survival of the largest individuals that emerged from the model. This reduction can also be introduced artificially to represent senescence or to impose an upper limit on survival rates that better represent our understanding of the real population (Needham et al., 2018). This approach can be particularly useful when modelling population dynamics of longevous trees, where we often lack data on mortality rates of the largest, often long-lived individuals (Clark et al., 2019; Xie et al., 2024). These individuals are typically difficult to observe in traditional census studies – although this may be improved by the use of large-scale remote sensing data.

### 2.6 | Analysing the IPM

A variety of life history traits that inform on the vitality and trajectory of a population can be easily calculated from stage structured population models (Salguero-Gómez et al., 2016), including IPMs (Bialic-Murphy et al., 2024; Needham et al., 2018). Some of these traits can be directly derived with only survival and growth information from the *P*-kernel (Zambrano & Salguero-Gómez, 2014). Multiple R packages exist for calculating these traits (*e.g.,* Jones et al., 2022; Stott et al., 2012; Stubben & Milligan, 2007). Here, we estimated the remaining life expectancy of the population using the *Rage* package (Jones et al., 2022). Life expectancy is the mean number of additional years that individuals (of a specified size) can be expected to live. Life expectancy has a direct impact on the biomass of the population (Needham et al., 2022), as well as being a proxy for carbon capture and storage, because large, long-lived trees will capture more carbon over their lifetimes than shorter-lived ones (Sobral et al., 2023).

We calculated remaining life expectancy across the full distribution of sizes, from the smallest trees – those entering the population at the lowest height limit (3 m) or minimum crown area (9 m^2^) – up to the largest observed individuals. Then, to illustrate how these models can help quantify the effect of a/biotic drivers on population dynamics, we tested the effect of increasing or decreasing topographic wetness index (TWI) and canopy density on life expectancy estimates (Fig. 3). Specifically, we compared mean conditions of TWI/canopy density with a scaled perturbation (*i.e.,* mean value ±50% of the variable’s standard deviation), to reflect typical variability in TWI and canopy density. To assess the impact of changing TWI and canopy density on life expectancy, we fit a linear model that included interactions between size class (minor or major), perturbation level (mean value, ±50%), and the variable being perturbed (TWI or canopy density). Additionally, we visually compared these model predictions to the range of possible life expectancy outcomes under different conditions of TWI/canopy density, up to the most extreme values of TWI/canopy density observed in this population.

**Figure 3.**
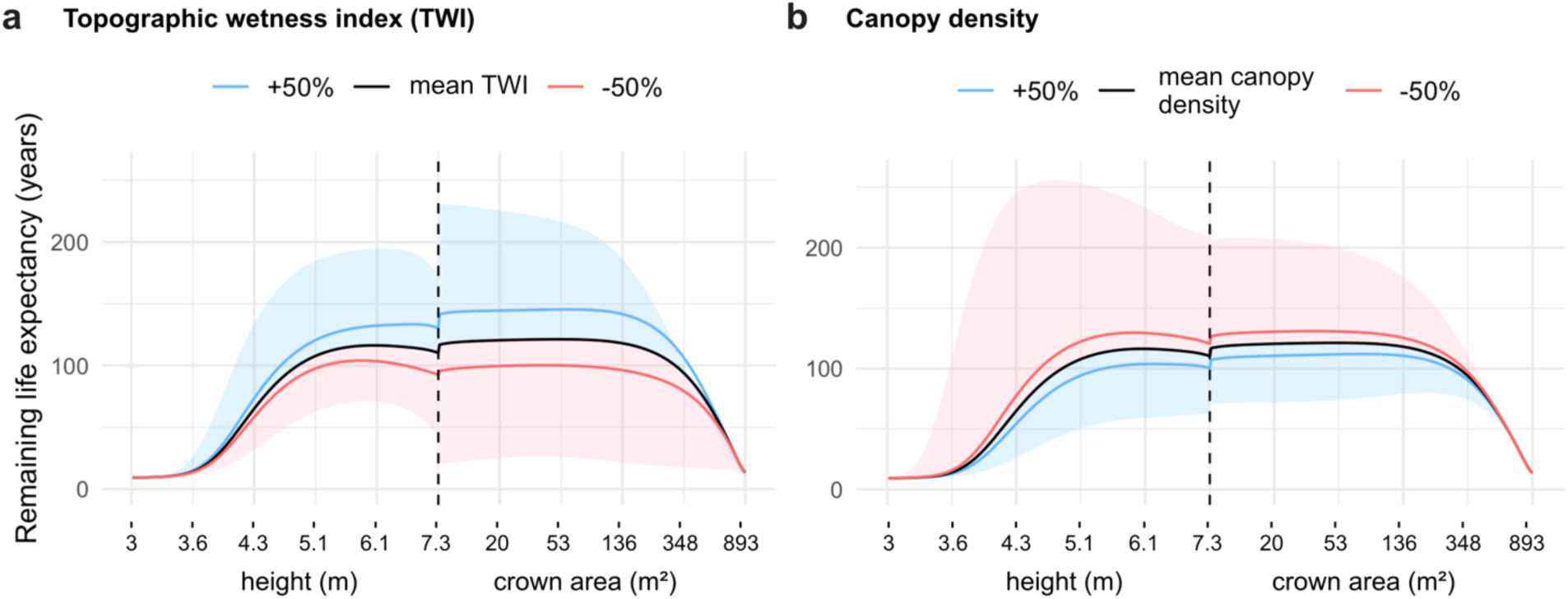
Effect of topographic wetness index (TWI) and canopy density on remaining life expectancy across tree sizes. The vertical dashed line divides trees into minor (left) and major (right) size classes. The black line represents life expectancy under mean conditions of TWI and canopy density observed in the 25km^2^ study site. Blue and red lines show life expectancy changes when TWI (**a**) or canopy density (**b**) are increased (blue) or decreased (red) by ±50% of the variable’s standard deviation. The perturbations are therefore scaled to reflect typical variability in TWI and canopy density. Shaded regions show the range of possible life expectancies under different conditions of TWI/canopy density, with upper and lower bounds representing the most extreme observed values in the site.

To further investigate the drivers of life expectancy, we examined the impact of perturbing individual parameters in each of the vital rate models (Eq. 1 & 2). In the supplementary materials, we provide an example of an elasticity analysis (Griffith, 2017), which can be used to assess the relative contribution of vital rate parameters to a life history trait of interest – in this case, mean life expectancy of the smallest individuals. While Figure 3 shows the relative impact of changing TWI and canopy density on remaining life expectancy, the elasticity analysis additionally reveals the demographic pathways through which these environmental factors may be driving a change in life expectancy (Easterling et al., 2000). For example, this pathway could be a reduction in survival rate of mature trees, or an absolute reduction in growth across size classes (Fig. S4).

## 3 | CASE STUDY RESULTS

We modelled the demographic changes of 42,213 Eucalypt trees using an integral projection model (IPM) parameterised with repeat LiDAR data. As a case study, we assessed the contribution of topographic wetness index (TWI) and canopy density to the life expectancy of trees across size classes. We found that remaining life expectancy was strongly linked to tree size, with trees at intermediate sizes (5.5-7.4 m in height, and 8.9-220 m^2^ in crown area) expected to have the highest number of additional years left to live (118 years on average; Fig. 3). Overall, large trees (those in the major class, see Fig. 2a) had a higher life expectancy (105 years) compared to smaller trees in the minor class (73 years), under average conditions of topographic wetness index and canopy density.

Changes to TWI (±50% of SD) had a larger impact on life expectancy than changes to canopy density. A 50% increase in TWI extended life expectancy by 26 years for major class trees (*p* < 0.001; Fig. 3a) and by 18 years for minor class trees (*p* = 0.235). Conversely, a 50% decrease in TWI led to a pronounced reduction in life expectancy, particularly for major class trees (−25 years; *p* < 0.001), while the reduction for minor class trees (−17 years) was non-significant (*p* = 0.265). Meanwhile, a 50% reduction in canopy density had a slight positive, albeit non-significant impact (Fig. 3b), increasing life expectancy by 7 years for both major class (*p* = 0.036) and minor class trees (*p* = 0.585).

## 4 | DISCUSSION

Forests are increasingly vulnerable to the impacts of climate change, disturbance, and land-use change (McDowell et al., 2020), and so effective monitoring is vital for protecting them and the ecosystem services they provide (Ohse et al., 2023). Unfortunately, ground-based forest surveys often fall short of meeting the data demands that demographic models require to build reliable forecasts, limiting their use across pertinent systems and scales (Doak et al., 2005). At the same time, vast amounts of remote sensing data are being generated (*e.g.* Kampe et al., 2010), yet remain under-utilised by population ecologists. To address this gap, we demonstrated a pipeline for parameterising a stage-structured model using repeat-LiDAR data. Using Australia’s Great Western Woodlands as a case study, we highlight the advantages and limitations of this approach, focussing on three areas that are key to broadening its application: 1) applying the pipeline to more complex systems, 2) capturing complete life histories, and 3) meeting data requirements for more robust demographic models.

### 4.1 | Challenges and opportunities in complex forests

Using a range of algorithms, it is now possible to automatically segment individual trees over large areas with reasonable accuracy (Battison et al., 2024; Wang et al., 2016). In this study, we segmented and modelled the vital rates of survival and growth for 42,213 individuals – far beyond what is typically available in demographic modelling studies (Levin et al., 2022; Salguero-Gómez et al., 2015). The eucalypt woodland featured a moderately sparse, open canopy, making it an ideal test system for our pipeline. However, the accuracy of tree segmentation is strongly tied to forest structure, with open canopies typically yielding better results than dense, multi-layered ones (Wang et al., 2016; Yang, Su, et al., 2019). The next challenge at the interface of repeat-LiDAR and ecological modelling lies in applying these pipelines to more complex systems, such as species-diverse forests with dense or multi-layered canopies. Tree segmentation algorithms trained on a single forest type may struggle to generalise to others without re-calibration, limiting their transferability (Wielgosz et al., 2024). Algorithms calibrated to a wide variety of forest types, or capable of handling diverse forest structures and data platforms are promising solutions (*e.g.,* deep-learning-based models; Henrich & Delden, 2024; Wielgosz et al., 2024). However, achieving more generalisable and user-friendly tree segmentation methods – requiring minimal tuning for specific use cases – will depend on access to publicly available, high-quality, labelled forest datasets.

Differentiating between species is also essential for building species-specific models that account for varying life histories and demographic processes. In this study, we focused on a functionally similar community of eucalypts and set a minimum size threshold to exclude smaller shrubs. While this simplified filtering approach was likely to be adequate for our study ecosystem, the study species do have different natural heights (*e.g.,* mature *Eucalyptus salubris* at the study site are typically 9-13 m whereas *Eucalyptus salmonophloia* are 10-20 m; Prober & Macfarlane, 2022) so might have different thresholds for switching from a dominance of height to crown growth. Similarly, more species-diverse forests may require more complex methods or integration of multiple data types. For example, hyperspectral remote sensing data has been used for continent-scale tree species classification, based on the variation in spectral properties between species (Marconi et al., 2022). Similarly, in a subtropical broadleaf forest, a combination of structural, spectral and textural features together (derived from LiDAR, hyperspectral, and RGB data) have been used to further enhance species classification accuracy (Qin et al., 2022). Although such approaches are not always viable due to cost or data availability, the development of multispectral-LiDAR systems could make this type of sensor fusion more feasible (Fassnacht et al., 2016; Takhtkeshha et al., 2024). Incorporating ground-based survey data will also be critical in addressing these challenges – for instance, by providing training data for species classification (Jia & Pang, 2023), or capturing important understory species that may be missed with airborne methods (Fassnacht et al., 2016).

### 4.2 | Completing the life cycle

Obtaining sufficient data on the vital rates of trees throughout their life cycle is a long-standing challenge in forest demography (Ohse et al., 2023; Ruiz-Benito et al., 2020). Here, we used repeat-LiDAR data to capture individual-level survival/mortality, growth, and dieback processes. A key advantage of LiDAR is its ability to capture changes in both crown size and tree height, making it more relevant than DBH for understanding the changing growth trajectories of trees (Laurans et al., 2024; Lines et al., 2022; Pretzsch et al., 2013). The results of our case study also indicate differential responses of small trees, which prioritise height growth, and large trees, which prioritise crown expansion, to biotic and abiotic factors such as canopy density and topographic wetness index. These findings underscore the importance of carefully selecting state variables that serve as ecologically meaningful vital rate predictors. However, by excluding trees smaller than 3 m – as a method for filtering out small shrubs, and as this was the minimum height of the manually delineated subset of tree crowns (Battison et al. 2024) – we overlooked the smallest and youngest trees in the population, which likely have distinct patterns of growth and survival (Canham & Murphy, 2017; Metcalf et al., 2009). Future demographic studies should aim to better capture these early life stages and shifting growth patterns across a wide range of tree species. This could be achieved through integration of LiDAR technologies (both airborne and terrestrial) with the extensive archive of field inventory data available through permanent forest plot networks that exist all over the world.

Mortality is particularly difficult to detect at the spatial scales and time intervals of most field surveys (McMahon et al., 2019). Here, we inferred mortality from threshold reductions in tree size or the disappearance of tree crowns, greatly improving our ability to detect tree deaths compared to traditional methods. However, LiDAR may miss mortality and dieback events in smaller trees, those obscured by dense canopies, or those at the early stages of decline. Our method may also misclassify trees as dead if they experience severe dieback within the survey period, but then later recover. Additional information, such as structural or spectral changes that precede death, could be used to improve mortality detection (Brodrick & Asner, 2017; Cotrozzi, 2022). Addressing these challenges is essential, as even small changes in mortality rates can have significant impacts on carbon cycling and ecosystem function (McMahon et al., 2019).

To make predictions about the long-term trajectory of populations, including growth rates, extinction risk, and carbon cycling, it is necessary to complete the life cycle by modelling reproductive output (Ohse et al., 2023). Building on the case study presented here, two fundamental components are needed at the very least to ‘close the loop’: 1) an estimate of how many individuals are recruited into the population between time steps, and 2) the size or age of trees that can produce seeds, to assign reproductive output (*e.g.,* using the “anonymous reproduction” method; Caswell, 2006). For recruitment, LiDAR could be used to track individuals entering a size class that can be reliably detected (*e.g.,* at least 3 m tall), which depends on forest density and point cloud resolution. By estimating the time taken for trees to reach this threshold size, it is then possible to infer the number of recruits produced by mature trees over time (Zambrano & Salguero-Gómez, 2014). To improve how recruits are attributed to mature trees, UAV or satellite imagery have been used to calculate flowering or fruiting probabilities at large spatial scales. For example, Lee et al. (2023) applied a deep learning model to a time series of UAV imagery to examine intraspecific variation in flowering, and Dixon et al. (2023) used multispectral satellite data (CubeSat daily images) to investigate flowering responses of eucalypts in relation to fire history. Additionally, experimental studies can provide more comprehensive measurements of reproductive output, such as seed production, germination rates, and seedling survival probabilities (Thomas, 2011). These field-derived estimates can be cross-validated with remote sensing data to calibrate model parameters, enabling more robust predictions of reproduction (Zipkin et al., 2021). Ultimately, collaborative work between these approaches and genetic analyses of parenthood (*e.g.,* Moran & Clark, 2012) will help us better understand individual sexual contributions to forest dynamics.

### 4.3 | The way forward for forest demographic modelling

While this pipeline is not an out-of-the-box solution for all forest systems and contexts, it provides a foundation for scaling demographic models with under-utilised LiDAR data. In the future, improving how these data are collected, processed, and integrated with other data types will help maximise its usefulness. For example, a key consideration is the choice of appropriate time intervals. If surveys do not span a sufficient period of time, they will not adequately capture growth of slow-growing trees (Munné-Bosch, 2020). On the other hand, surveys conducted too far apart risk missing events such as recruits that establish and die between observations. The ideal balance would be a greater number of surveys conducted at more frequent intervals. Open data initiatives, such as NEON (Kampe et al., 2010), and more affordable platforms, like UAVs, are making this more achievable (Araujo et al., 2021; Kleinsmann et al., 2023). The additional context gained from more frequent surveys (*e.g.,* leaf-on and leaf-off conditions) can also be used to improve segmentation or species classification algorithms (Chen et al., 2022).

Another challenge lies in the comparability of LiDAR surveys. For instance, potential issues arise when the data acquisition parameters vary greatly between flights (*e.g.,* pulse density, flight elevation; Næsset, 2009), or in the way the data are processed (*e.g.,* to produce canopy height models). Applying a standardised processing pipeline to all LiDAR point clouds in the analysis can mitigate these issues (Fischer et al., 2024). Using algorithms that perform well when applied to data of varying quality can also improve the reliability of results (Wielgosz et al., 2024). Additionally, certain metrics, such as those derived from canopy height models, rather than directly from 3D point clouds, tend to be more robust to these differences, aiding comparisons across time and space (Fischer et al., 2024; Zhang et al., 2024).

High-quality, standardised data from more frequent LiDAR surveys will greatly benefit our ability to build ecologically informative demographic models. While field surveys excel at species identification, understanding reproductive strategies, and capturing understory dynamics (Obeso, 2002), remote sensing offers the scalability needed to track forest responses to climate and disturbance at broad spatial and temporal scales. By leveraging the strengths of both approaches, we can build models that are better equipped to inform conservation and management strategies in a rapidly changing world.

## Acknowledgements

We acknowledge and pay our respects to the First Nations people in the Great Western Woodlands on whose land this work was conducted. The project was funded by a Western Australian State NRM grant to the Ngadju Conservation Aboriginal Corporation and CSIRO and supported by the Australian Government through the TERN Great Western Woodlands Supersite and the TERN Landscapes platform. Airborne LiDAR from 2021 was collected by Landgate (Western Australia’s land information authority) under the Capture WA program with additional support from the Western Australian Department of Biodiversity, Conservation and Attractions Goldfields Region office. AR was supported by a BBSRC DTP studentship. CMH and RSG were supported by a NERC Pushing the Frontiers grant to RSG (NE/X013766/1).

## Author contributions

AR and RSG developed the ideas, with contributions from TJ. Lidar data were processed and segmented by RB. JM, OS, SM, CMH, and SM contributed to the interpretation of IPM code developed by RSG and AR. SP contributed to the ecological interpretation of results in the context of the study site. AR wrote the first version of the manuscript with input from RSG. All coauthors contributed to the final version of this manuscript.

