## Supplementary Materials for "Modelling forest dynamics using integral projection models (IPMs) and repeat LiDAR"

**Contents**

1. Point cloud processing: producing CHMs and DTMs 2
2. Primary and secondary growth of eucalypts 3
3. Choosing a suitable cut-off point between two continuous traits 4
4. Sensitivity of remaining life expectancy to the choice of cut-off 6
5. Model comparison AIC table 8
6. Elasticity analysis 10
7. References 12

### 1. Point cloud processing: producing CHMs and DTMs

Georeferenced point clouds for both airborne laser scanning (ALS) surveys (2012 and 2021) were processed using a combination of CloudCompare (https://danielgm.net/cc), QGIS (https://qgis.org) and R version 4.1.0 (R Core Team, 2021). First, outliers and duplicate points were removed using a statistical outlier removal filter. Then, the point clouds were classified into ground and non-ground returns, with the former being used to create a digital elevation model (DEM) for each survey. To produce the canopy height model (CHM), the elevations of non-ground points were then subtracted from the DEM to produce a 0.5 m resolution normalised CHM using a pit-free algorithm in the *lidR* R package (Khosravipour et al., 2014; Roussel et al., 2020). The two CHMs (2012 and 2021) were aligned and clipped to retain only areas of overlap, leaving 24.98 km$^2$. Additionally, to explore the influence of topographic factors on tree vital rates, a 5 m resolution digital terrain model (DTM) was created from the 2021 ground returns using the triangular irregular network (TIN) algorithm from the *lidR* package.

### 2. Primary and secondary growth of eucalypts

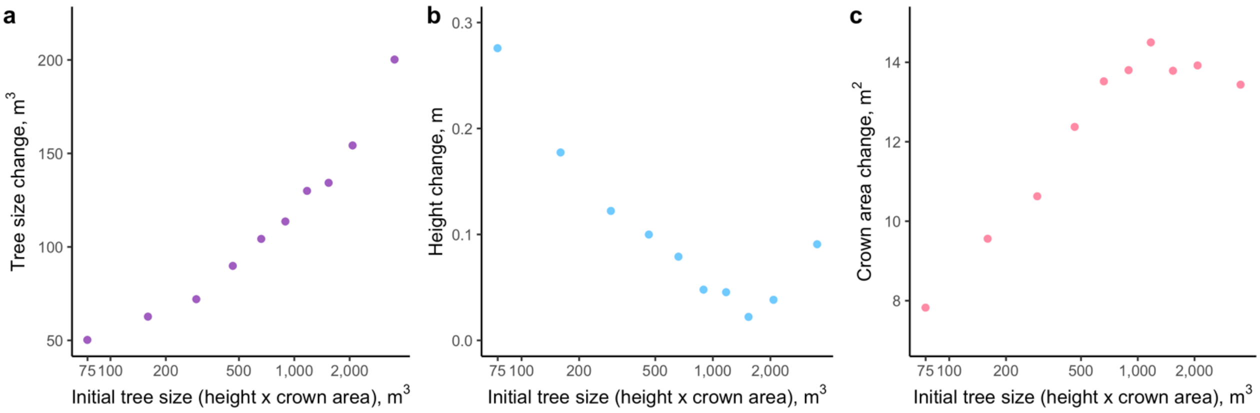


**Figure S1.** Change in direction of growth: eucalypt trees initially prioritise growing taller, before expanding their crowns. To visualise the changing growth trajectory, individual tree crowns were split into 10 equal-sized bins based on initial tree size (the product of height and crown area in 2012). **a**) Overall tree size change increases with initial size, but this trait does not reveal the different ways that trees are allocating resources to growth when they are small compared to when they are large. **b**) On average, smaller trees experienced greater height growth than large trees, meanwhile, larger trees experienced more crown area growth than smaller ones (**c**). Using a two-stage model, as in this case study, therefore allows us to consider these processes separately.

### 3. Choosing a suitable cut-off point between two continuous traits

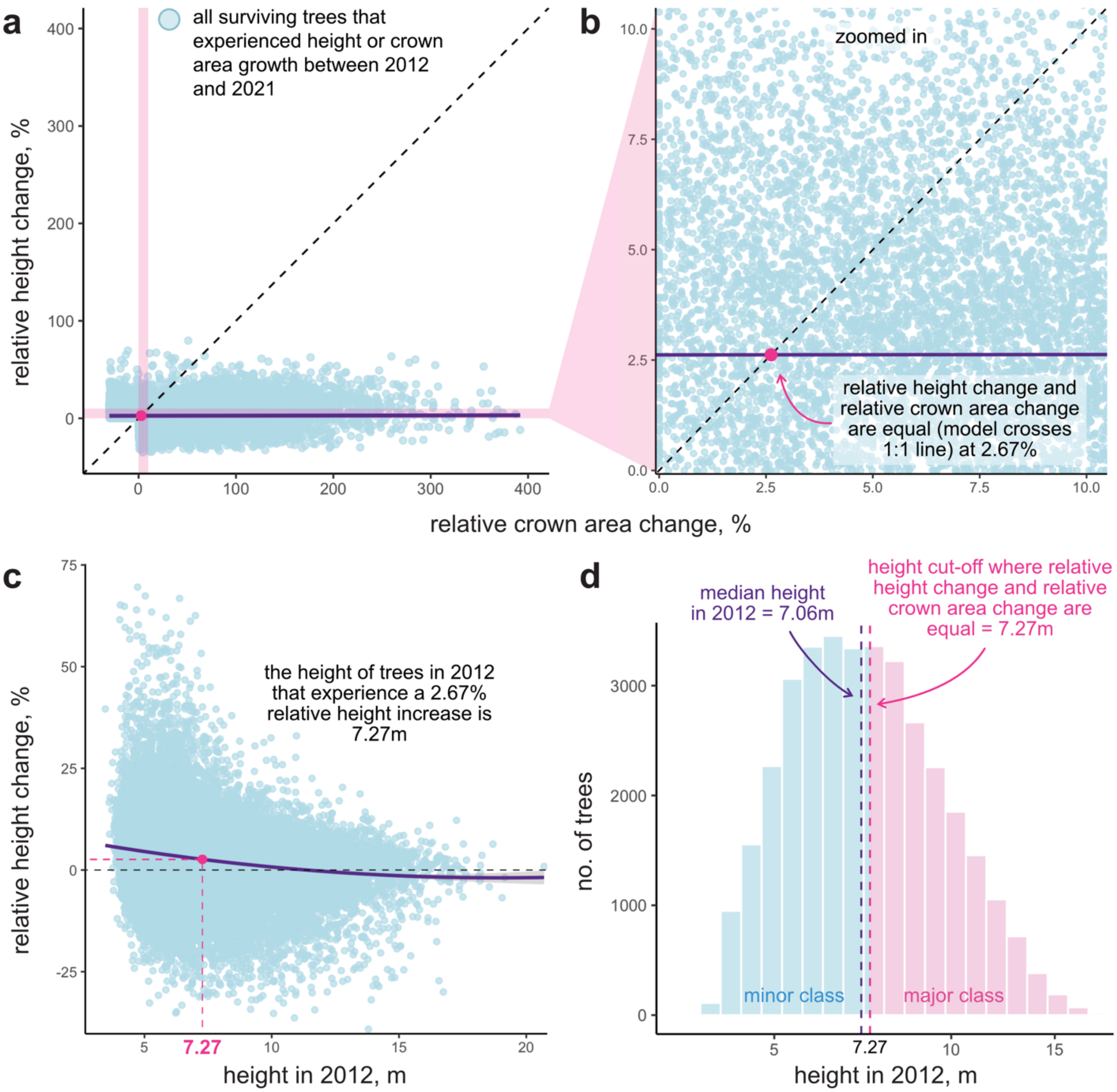


**Figure S2**. Process for determining a suitable cut-off point between small trees (minor class) and large trees (major class), for construction of a two-stage integral projection model (IPM). **a**) Relative height change (%) *versus* relative crown area change (%) of all surviving individual trees that experienced at least one form of growth (i.e., height growth or crown area growth) between 2012 and 2021. Dashed line indicates the 1:1 line, where relative height and crown area change are equal. The purple line is a linear model describing the relationship between relative height change and relative crown area change. **b**) is a zoomed in version of panel **a**, where only the datapoints at the intersection of the pink highlighted regions are shown. The pink dot shows the point at which the model (purple line) crosses the 1:1 line. This is the point at which relative height change and relative crown area change are predicted to be equal. **c**) 7.27 m is the height of trees in 2012 that are predicted to experience an equal relative height and crown area increase from $t$ to $t$+1. **d**) Histogram of tree heights in 2012 (log scale with back-transformed value labels). The chosen cut-off (7.27 m) is close to the median height value in 2012 (7.06 m).

4. **Sensitivity of remaining life expectancy to the choice of cut-off in this two-stage IPM**

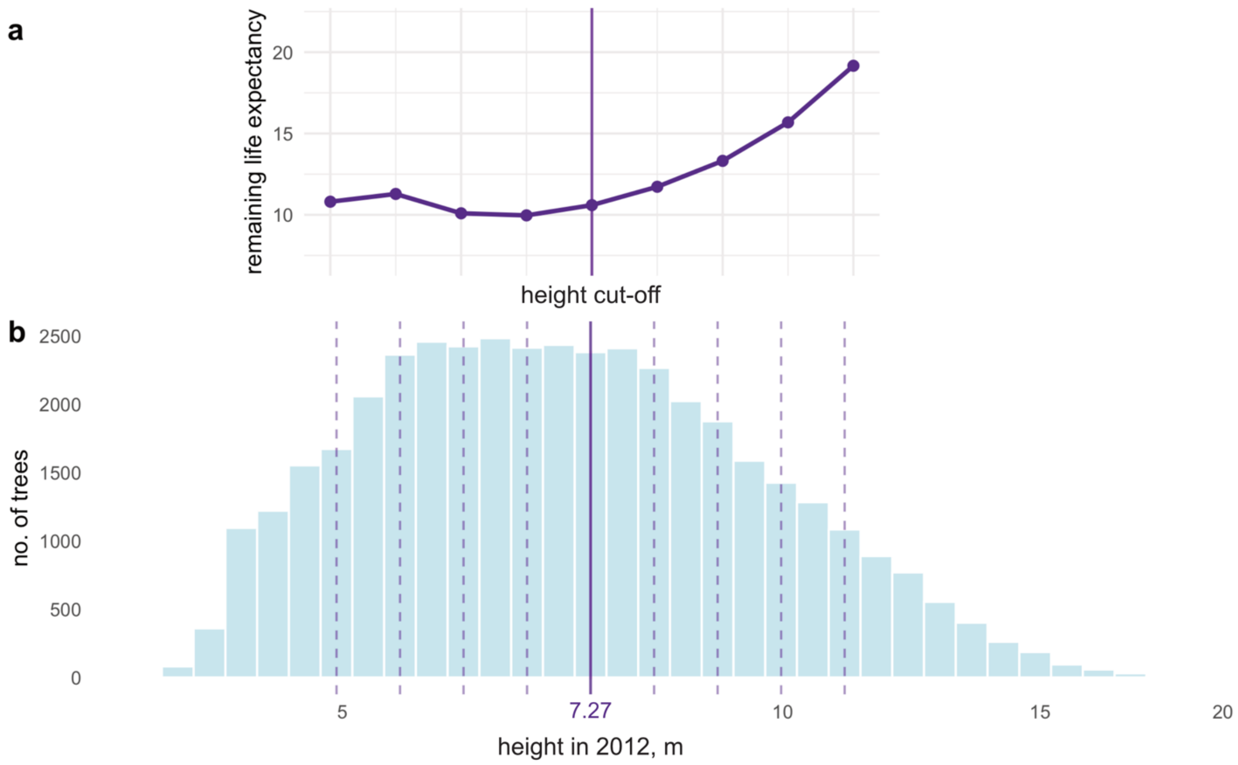


**Figure S3**. Sensitivity of remaining life expectancy to the choice of cut-off. **a**) Life expectancies were calculated using the *Rage* R package (Jones et al., 2022). In this sensitivity analysis, life expectancy refers to the mean number of additional years that the smallest individual trees included in the IPM can expect to live, *i.e.*, trees 3 m to 3.6 m in height (see Fig. S4). The estimate corresponds to individuals represented in the first 10% of bins in the IPM, which were given equal weighting in the life expectancy calculation. **b**) Distribution of tree heights in 2012 (log-scale, with back transformed x-axis labels). The solid purple vertical line represents the actual cut-off value used to construct the sub-kernels of the IPM, while dashed purple vertical lines indicate alternative cut-off values tested in this brute-force sensitivity analysis.

Note of caution when constructing a two (or more) stage IPM based on different traits (e.g., height and crown area): There may be a step-change in survival rate around the cut-off point between the two traits, because the intercept of the survival function will likely change from one trait to the next. This survival function directly influences the life expectancy estimates, sometimes resulting in a step-change in life expectancy at the corresponding stage of this cut-off (Fig 3). Therefore, when calculating life expectancy (or other life history traits) for individuals positioned around this cut-off point, it is important to be aware of the impact this may have on the sensitivity of a life history trait estimate in this scenario. In other words, small perturbations in vital rate parameters, or even small adjustments to the choice of cut-off, can have very large impacts on the life expectancy of these individuals due to this step-change. To minimise this sensitivity issue, one could use a mixing distribution that does not give too much weighting to the individuals in those problematic stages immediately before and after the cut-off.

### 5. Model comparison AIC table

**Table S1.** Heights, crown areas, TWI and canopy density are log-transformed

| **model** | AIC | delta AIC | akaike weight |
|---|---|---|---|
| minor class survival | | | |
| survival ~ height 2012 | 14718 | 408 | 0 |
| survival ~ height 2012 (quadratic) | 14442 | 123 | 0 |
| survival ~ height 2012 + TWI | 14711 | 401 | 0 |
| survival ~ height 2012 (quadratic) + TWI | 14433 | 123 | 0 |
| survival ~ height 2012 + TWI + canopy density | 14555 | 245 | 0 |
| **survival ~ height (quadratic) + TWI + canopy density** | **14310** | **0** | **1** |
| minor class remain | | | |
| remain ~ height 2012 | 8892 | 197 | 0 |
| remain ~ height 2012 (quadratic) | 8839 | 143 | 0 |
| remain ~ height 2012 + TWI | 8798 | 103 | 0 |
| remain ~ height 2012 (quadratic) + TWI | 8743 | 48 | 0 |
| remain ~ height 2012+ TWI + canopy density | 8753 | 58 | 0 |
| **remain ~ height (quadratic) + TWI + canopy density** | **8695** | **0** | **1** |
| minor class growth | | | |
| height 2021 ~ height 2012 | -42582 | 249 | 0 |
| height 2021 ~ height 2012 (quadratic) | -42720 | 111 | 0 |
| height 2021 ~ height 2012 + TWI | -42685 | 146 | 0 |
| height 2021 ~ height 2012 (quadratic) + TWI | -42827 | 4 | 0.13 |
| height 2021 ~ height 2012 + TWI + canopy density | -42693 | 138 | 0 |
| **height 2021 ~ height 2012 (quadratic) + TWI + canopy density** | **-42831** | **0** | **0.87** |
| minor class growth (residuals) | | | |
| residuals ~ 1 | -57184 | 40 | 0 |
| residuals ~ height 2012 | -57198 | 27 | 0 |
| **residuals ~ height 2012 (quadratic)** | **-57225** | **0** | **1** |
| promotion class growth | | | |
| crown area 2021 ~ height 2012 | 4385 | 377 | 0 |
| crown area 2021 ~ height 2012 (quadratic) | 4364 | 356 | 0 |
| crown area 2021 ~ height 2012 + TWI | 4385 | 377 | 0 |
| crown area 2021 ~ height 2012 (quadratic) + TWI | 4364 | 356 | 0 |
| crown area 2021 ~ height 2012 + TWI + canopy density | 4023 | 15 | 0 |
| **crown area 2021 ~ height 2012 (quadratic) + TWI + canopy density** | **4008** | **0** | **1** |
| promotion class growth (residuals) | | | |
| residuals ~ 1 | 1880 | 13 | 0 |

| | | | |
|---|---|---|---|
| residuals ~ height 2012 | 1875 | 7 | 0.02 |
| **residuals ~ height 2012 (quadratic)** | **1867** | **0** | **0.97** |
| major class survival | | | |
| survival ~ crown area 2012 | 9449 | 283 | 0 |
| survival ~ crown area 2012 (quadratic) | 9450 | 285 | 0 |
| survival ~ crown area 2012 + TWI | 9190 | 24 | 0 |
| survival ~ crown area 2012 (quadratic) + TWI | 9189 | 24 | 0 |
| survival ~ crown area 2012 + TWI + canopy density | 9166 | 0 | 0.46 |
| **survival ~ crown area 2012 (quadratic) + TWI + canopy density** | **9165** | **0** | **0.54** |
| major class remain | | | |
| remain ~ crown area 2012 | 7885 | 172 | 0 |
| remain ~ crown area 2012 (quadratic) | 7764 | 51 | 0 |
| remain ~ crown area 2012 + TWI | 7886 | 173 | 0 |
| remain ~ crown area 2012 (quadratic) + TWI | 7766 | 53 | 0 |
| remain ~ crown area 2012 + TWI + canopy density | 7826 | 113 | 0 |
| **remain ~ crown area 2012 (quadratic) + TWI + canopy density** | **7713** | **0** | **1** |
| major class growth | | | |
| crown area 2021 ~ crown area 2012 | -13801 | 720 | 0 |
| crown area 2021 ~ crown area 2012 (quadratic) | -14455 | 65 | 0 |
| crown area 2021 ~ crown area 2012 + TWI | -13887 | 633 | 0 |
| crown area 2021 ~ crown area 2012 (quadratic) + TWI | -14511 | 9 | 0.01 |
| crown area 2021 ~ crown area 2012 + TWI + canopy density | -13899 | 621 | 0 |
| **crown area 2021 ~ c. area 2012 (quadratic) + TWI + canopy density** | **-14520** | **0** | **0.99** |
| major class growth (residuals) | | | |
| residuals ~ 1 | -26545 | 949 | 0 |
| residuals ~ crown area 2012 | -27326 | 167 | 0 |
| **residuals ~ crown area 2012 (quadratic)** | **-27494** | **0** | **1** |
| demotion class growth | | | |
| height 2021 ~ crown area 2012 | -2856 | 1 | 0.16 |
| height 2021 ~ crown area 2012 (quadratic) | -2855 | 2 | 0.11 |
| height 2021 ~ crown area 2012 + TWI | -2856 | 1 | 0.15 |
| height 2021 ~ crown area 2012 (quadratic) + TWI | -2855 | 2 | 0.11 |
| **height 2021 ~ crown area 2012 + TWI + canopy density** | **-2857** | **0** | **0.29** |
| height 2021 ~ crown area 2012 (quadratic) + TWI + canopy density | -2856 | 1 | 0.19 |
| demotion class growth (residuals) | | | |
| residuals ~ 1 | -3815 | 7 | 0.03 |
| **residuals ~ crown area 2012** | **-3821** | **0** | **0.97** |

### 8. Elasticity analysis

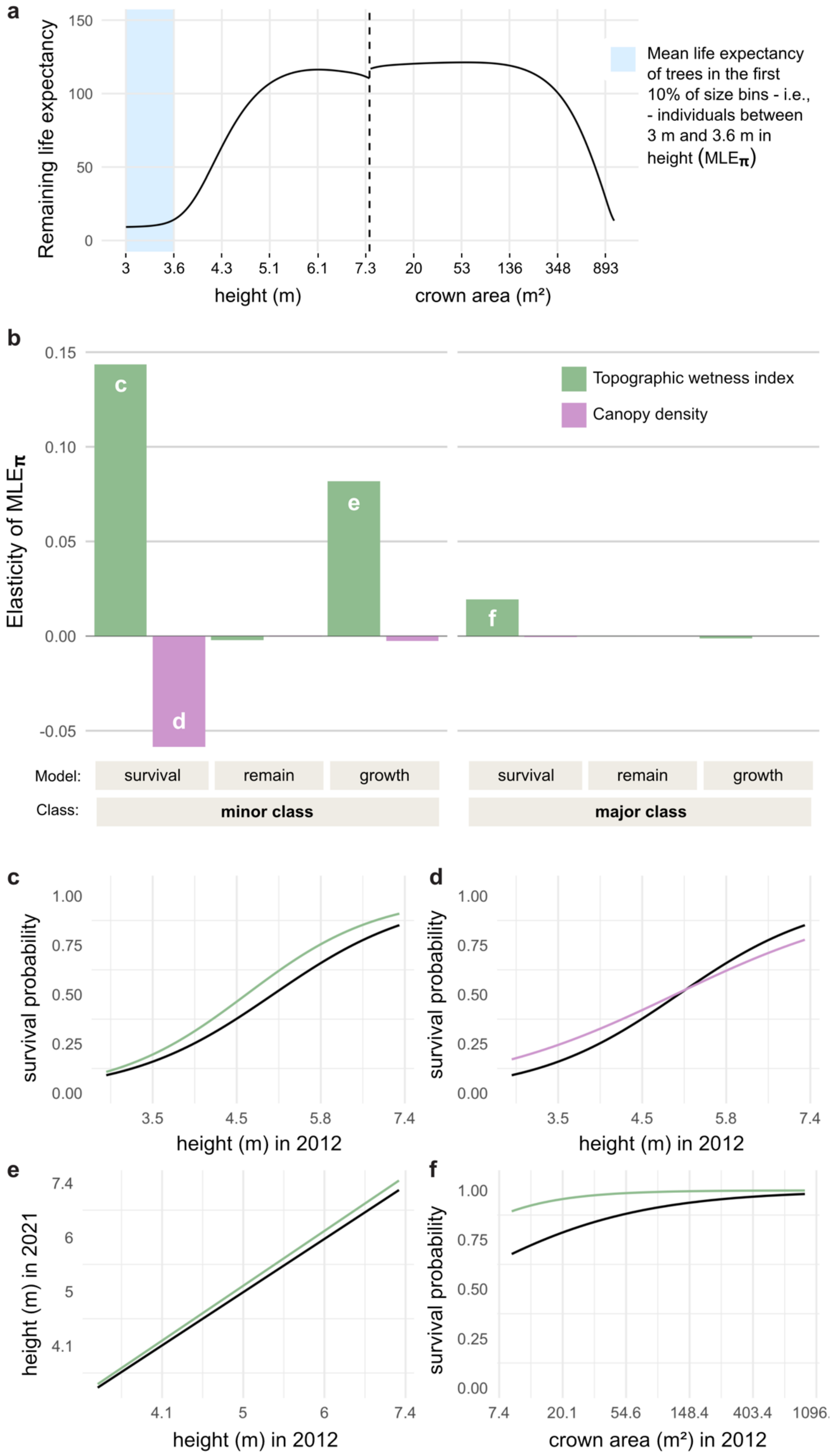

**Figure S4**. Elasticity analysis, *i.e.,* the proportional change in life expectancy that results from a proportional change in each of the vital rate parameters **a**) In this elasticity analysis, the life history trait of interest is the mean life expectancy of the smallest individuals, *i.e*., trees 3 m to 3.6 m in height. This life expectancy estimate (hereafter referred to as $\mathrm{MLE}_{\pi}$) corresponds to individuals represented in the first 10% of bins, which were given equal weighting in the life expectancy calculation. **b**) The relative contribution of vital rate parameters to $\mathrm{MLE}_{\pi}$. Parameters relating to topographic wetness index (TWI; green) and canopy density (purple) for each of the minor class and major class vital rate functions are shown. Letters inside the bars correspond to panels **c** – **f**. In this case study, $\mathrm{MLE}_{\pi}$ is most elastic to the survival of minor class trees, particularly through a change in topographic wetness index. Panels **c** – **f** show how the parameter perturbations in panel **b** impact the shape of the vital rate functions. For instance, an increase in TWI shifts the survival probability upwards for all trees in the minor class (**c**). However, the mechanism in **d** is more complex: survival probability is increased for individuals up to 5.15 m in height, while survival probability is reduced for individuals larger than this. Overall, the elasticity analysis reveals that $\mathrm{MLE}_{\pi}$ is more sensitive to changes in TWI than to changes in canopy density, and TWI has a generally positive impact on $\mathrm{MLE}_{\pi}$, while canopy density has a smaller, negative impact on $\mathrm{MLE}_{\pi}$.
